# Preclinical screen for protection efficacy of chlamydial antigens that are immunogenic in humans

**DOI:** 10.1101/2023.08.31.555742

**Authors:** Chunxue Lu, Jie Wang, Guangming Zhong

## Abstract

**To search for subunit vaccine candidates, immunogenic chlamydial antigens identified in humans were evaluated for protection against both infection and pathology in a mouse genital tract infection model under three different immunization regimens. The intramuscular immunization regimen was first used to evaluate 106 chlamydial antigens, which revealed that two antigens significantly reduced while 11 increased genital chlamydial burden. The two infection-reducing antigens failed to prevent pathology and 23 additional antigens even exacerbated pathology. Thus, intranasal mucosal immunization was tested next since intranasal inoculation with live *C. muridarum* prevented both genital infection and pathology. Two of 29 chlamydial antigens evaluated were found to prevent genital infection but not pathology and three exacerbate pathology. To further improve protection efficacy, a combinational regimen (intranasal priming + intramuscular boosting + a 3^rd^ intraperitoneal/subcutaneous boost) was tested. This regimen identified 4 infection-reducing antigens but only one of them prevented pathology. Unfortunately, this protective antigen was not advanced further due to its amino acid sequence homology with several human molecules. Two pathology-exacerbating antigens were also found. Nevertheless, intranasal mucosal priming with viable *C. muridarum* in control groups consistently prevented both genital infection and pathology regardless of the subsequent boosters. Thus, screening 140 different chlamydial antigens with 21 repeated multiple times in 17 independent experiments failed to identify a subunit vaccine candidate but the efforts have revealed pathogenic antigens and demonstrated the superiority of viable chlamydial organisms in inducing immunity against both genital infection and pathology, laying the foundation for developing an attenuated live Chlamydia vaccine.**

**Importance:** **This manuscript describes a systematical effort in searching for a chlamydial subunit vaccine by taking advantage of both the immunogenic chlamydial antigens identified in humans and a robust mouse genital tract infection model for simultaneously evaluating protection against both genital infection and pathology. Screening 140 different chlamydial antigens (21 repeated multiple times) using three different immunization regimens in 17 independent experiments identified no subunit vaccine candidate. Nevertheless, the efforts revealed multiple pathogenic chlamydial antigens and demonstrated the superiority of mucosal inoculation with viable chlamydial organisms for inducing immunity against both genital infection and pathology, suggesting that a live attenuated Chlamydia vaccine strategy should be considered.**

## Introduction

*Chlamydia trachomatis* is a leading cause of sexually transmitted infection in the human genital tract, which, if not treated in time, may lead to pathological sequelae in the upper genital tract such as hydrosalpinx, resulting in loss of reproductive function (1-4). Although the precise mechanism remains unclear, invading the genital tract epithelial cells is considered a key determinant in chlamydial pathogenicity (5-7). Chlamydia mainly targets epithelial cells for infection, starting with the entry by an infectious elementary body (EB). The internalized EB differentiates into a noninfectious but metabolically active reticulate body (RB) that replicates, during which, new molecules are synthesized for secretion outside of the chlamydial organisms (8-13) and metabolites (14) are generated that may also be released into host cells. After completing the replication, the progeny RBs differentiate back into EBs for spreading to neighboring host cells. It is worth noting that chlamydial secretion molecules and/or metabolites may not become part of the chlamydial organisms but can interact with host cells, impacting both chlamydial pathogenicity (15-18) and host responses (19, 20). It will be interesting to determine whether the chlamydial infection-dependent induction of host responses is necessary for preventing subsequent chlamydial infection and pathogenicity.

Although effective antibiotics are available for treating chlamydial infection, the challenge is that *C. trachomatis* infection is often asymptomatic or lacks specific symptoms, thus significantly hindering the treatment efficacy. Obviously, vaccination offers a long-term solution to this challenge. However, despite the extensive efforts in more than half-century (21-39), there is still no licensed Chlamydia vaccine for humans. The *C. muridarum* intravaginal infection mouse model has been extensively used to study *C. trachomatis* pathogenesis and immunology (19, 40-46). Although *C. muridarum* causes no known diseases in humans, it can infect the female mouse genital tract productively (41) and following a single intravaginal inoculation, *C. muridarum* induces many different strains of mice to develop upper genital tract pathologies such as hydrosalpinx that closely resembles that in the human genital tracts induced by *C. trachomatis* (42, 47). Thus, this mouse genital tract infection model has been extensively used to search for chlamydial vaccine candidates. Since our whole genome-scale soluble protein-based proteome array analyses of sera from *C. trachomatis*-infected women had systemically identified chlamydial immunogenic antigens in humans (3, 48, 49), it was logical to use the murine model to evaluate the protection efficacy of the chlamydial antigens recognized by humans.

The current manuscript reports a portion of the results from our past vaccine research efforts. The goal of these experiments was to search for subunit vaccine antigens using the *C. muridarum* infection of mouse genital tract model. To maximize the protection efficacy induced by chlamydial antigens, the *C. muridarum* homologues of the *C. trachomatis* antigens were used as the immunogens in most cases. Mice were immunized 3 times with chlamydial antigens and one month after the last immunization, mice were intravaginally challenged with *C. muridarum* and monitored for genital tract chlamydial burden and pathology. Three different vaccination regimens were compared. Chlamydial antigens adjuvanted with CpG and incomplete Freund’s adjuvant (IFA) were used for intramuscular (IM), intraperitoneal (IP) or subcutaneous (SC) immunization while those adjuvanted with CpG alone for intranasal (IN) mucosal inoculation. The exception was that low doses of live *C. muridarum* were intranasally inoculated without any adjuvant. An intramuscular immunization regimen was first used to screen a total of 106 chlamydial antigens, which revealed two antigens that significantly reduced genital chlamydial burden while surprisingly 11 increased the burden comparing to the negative immunization controls. The 2 infection-reducing antigens are recombinant protein TC0420 and native major outer membrane protein (MOMP) extracted from *C. muridarum* and their protection efficacies were as potent as that induced by adjuvanted UV-inactivated *C. muridarum* but significantly less potent than that induced by adjuvanted live *C. muridarum*. Unfortunately, the intramuscular immunization-induced immunity failed to reproducibly prevent genital pathology regardless of the immunogens used. Further, under the intramuscular immunization regimen, 23 antigens were found to significantly exacerbate genital pathology, raising a concern on the safety of this regimen. Thus, an intranasal mucosal immunization regimen was tested next since intranasal inoculation with live *C. muridarum* consistently protected against both genital tract infection and pathology although UV-inactivated *C. muridarum* failed to do so (50). Two of 29 chlamydial antigens evaluated were found to be protective against genital infection but not pathology. Further, three antigens were found to exacerbate pathology. To further boost the immunogenicity of chlamydial antigens, a combinational regimen (intranasal priming + intramuscular boosting + a final intraperitoneal or subcutaneous booster) was tested, which identified 4 antigens that significantly prevented genital infection with only one of them also preventing pathology. However, the antigen that induced protection against both genital infection and pathology was not recommended to move forward further due to its amino acid sequence homology with human molecules. In addition, two other antigens were also found to exacerbate pathology under this regimen. Nevertheless, in the control groups, as long as mice were intranasally primed with live *C. muridarum*, protection against both genital chlamydial infection and pathology was always induced regardless of the subsequent boosters, which has again demonstrated the dependence of inducing immunity against both genital infection and pathology on viable chlamydial organisms. Thus, although screening 140 different chlamydial antigens with 21 of them repeated multiple times in 17 independent experiments failed to identify a subunit chlamydial vaccine antigen for humans, the extensive efforts have revealed multiple pathogenic antigens and more importantly demonstrated the superiority of viable chlamydial organisms in priming protective responses against both genital infection and pathology. Our findings suggest that an attenuated live Chlamydia vaccine strategy should be considered given both the past success in using similar strategies for developing human vaccines (51-53) and the recent progresses made in this direction (54-59).

## Materials and Methods

### Chlamydial vaccine antigen expression and purification

The chlamydial vaccine antigens were identified by using a whole genome-scale soluble GST (glutathione-s-transferase) fusion protein array consisting of 908 distinct *C. trachomatis* antigens to screen sera from 99 women infected with *C. trachomatis* (48). A total of 27 antigens were found to react with 50% or more patient sera, which were designated as immunodominant antigens, and 719 antigens were recognized by at least one patient serum, which were designated as immunogenic antigens. The most immunogenic chlamydial antigen identified in humans was the *C. trachomatis* serovar D plasmid-encoded glycoprotein 3 (Pgp3 encoded by pCT03, its homologue in *C. muridarum* is encoded by pCM03) that was recognized by 96% patients followed by the chlamydial protease/proteosome-like activity factor (CPAF, encoded by ORF CT858, its homologue in *C. muridarum* is encoded by TC0248) recognized by 94% and the outer membrane complex protein B (OmcB by ORF CT443, its homologue in *C. muridarum* is encoded by TC0727) by 89%. The major outer membrane protein (MOMP encoded by ORF CT681, its homologue in *C. muridarum* is encoded by TC0052) was ranked the 22^nd^ since it was recognized by only 58% patient sera. A total of 140 different chlamydial antigens (not counting repeating or antigens in different versions or with different fusion partners) from the top of the immunogenic antigen list were selected for protection efficacy evaluation in a *C. muridarum* infection of mouse genital tract model (see Table 1S for a list of all antigens tested). To maximize the induction of protection efficacy in the *C. muridarum*-mouse model, the corresponding *C. muridarum* homologues were used as immunogens. The open reading frames (ORFs) coding for each of the 140 *C. muridarum* antigens were cloned from the *C. muridarum* genome and plasmid (NCBI protein accession# AAF39550, http://www.ncbi.nlm.nih.gov/sites/entrez; ref: (60)) into pGEX vector (Amersham Pharmacia Biotech, Inc., Piscataway, NJ) and expressed as fusion proteins with glutathione-s-transferase (GST) fused to the N-terminus of the chlamydial proteins. The resultant fusion proteins were purified using a glutathione-conjugated agarose beads as described previously (61-63). In most cases, the full-length chlamydial antigens were purified and used as immunogens. When it was difficulty in expressing an antigen in full-length, only a fragment of that antigen was expressed instead. The C-terminal region was indicated with a “c”, N-terminus with “n”, middle region with “m” while “s” was used to indicate the short form of Pgp3 as described previously (64, 65). For some experiments, two antigens were fused together as a bi-valent vaccine antigen or for manipulating the conformation of a target antigen/fragment, including the fusion of the full-length *C. muridarum* MOMP with the C-terminus of pCM03 (TC0052-pCM03c) and the MOMP C- and N-terminal regions with Pgp3c (TC0052c-pCM03c & TC0052n-pCM03c) respectively. The N- and C-terminal regions of OmcB were also fused to CPAFc and CM Pgp3c (TC0727n-TC0248c & TC0727c-pCM03c) respectively. In most cases, chlamydial antigens were purified by cleaving from the beads and the soluble proteins were concentrated via centrifugal concentrators. When cleavage was difficulty, the GST-chlamydial fusion proteins were eluted off the beads and used as immunogens. Free GST was eluted as negative immunization control antigen. In some cases, the elution of fusion protein was difficulty, the bead complexes after extensive washing were then used as immunogens for intramuscular or intraperitoneal injection.

For positive immunization controls, live *Chlamydia muridarum* (Nigg3 strain) EBs were used. The organisms were grown in HeLa cells (ATCC, Manassas, VA 20108), purified and titrated as described previously (66, 67). To prepare UV-inactivated *C. muridarum* EBs (UV-EB or UV-CM), EBs were exposed to UV light from a G36T5L/C UV lamp (Universal light source, San Francisco, CA) at 5cm for 45 min at room temperature. To ensure that the UV-CM organisms were completely inactivated, each UV-EB preparation was tested on HeLa cells and no inclusion-forming unit (IFU) was recovered after incubation for 24h (data not shown). Aliquots of both live CM and UV-CM organisms were stored at -80°C till use.

### Mouse immunization

Female Balb/c mice were purchased at the age of 4 to 5 weeks old from Charles River Laboratories, Inc. (Wilmington, MA). For intramuscular, intraperitoneal, or subcutaneous immunization, 30μg of each chlamydial antigen or GST protein or 1 × 10^5^ IFUs of UV-CM or live CM plus 10μg CpG in a total volume of 50μl PBS emulsified in equal volume of IFA (incomplete Freund’s Adjuvant) was used for each injection. The CpG with a sequence of 5’-TCC.ATG.ACG.TTC.CTG.ACG.TT-3’ (all nucleotides are phosphorothioate-modified at the 3’ inter-nucleotide linkage) was purchased from Integrated DNA Technologies (IDT, Coralville, IA) and the IFA from Sigma-Aldrich (St. Louis, MO). We used the CpG-IFA as adjuvants for the intramuscular injection because they have been shown to induce Th1 dominant immune responses (68, 69). For intranasal mucosal immunization, 30μg of chlamydial antigen or GST protein or 1 × 10^6^ IFUs of UV-CM plus 10 μg CpG or 50 or 500 IFUs of live CM alone in a total volume of 20μl PBS was used for each inoculation. In some experiments, DDA (dimetyldiocta-decylammonium bromide [product no. 890810P]) and TDB [D-(+)- trehalose 6,6’-dibehenate (product no. 890808P), both from Avanti Polar Lipids (Alabaster, AL)] were used since DDA/TDB were reported to promote immunogenicity of chlamydial antigens (70). The procedure for formulating DDA/TDB at a final inoculation dose of 250μg DDA and 50μg TDB in a total immunization volume of 200 μl per injection per mouse was described previously (70).

Three different immunization regimens were compared in the current study: Intramuscular, intranasal, and combinational regimens. For all regimens, mice were immunized 3 times, on day 0, day 20 and day 30 respectively. The exception is the intranasal immunization with live CM, which was given only once at the beginning of the immunization schedule. The combinational regimen consists of an intranasal priming and an intramuscular boosting with or without a final intraperitoneal or subcutaneous booster.

### Mouse challenge infection

Thirty days after the final immunization, each mouse was inoculated intravaginally with 2 × 10^4^ IFUs of live *C. muridarum* organisms in 20μl of SPG (sucrose-phosphate-glutamate buffer consisting of 218mM sucrose, 3.76mM KH2PO4, 7.1mM K2HPO4, 4.9mM glutamate, pH 7.2). Five days prior to the inoculation, each mouse was injected with 2.5mg Depo-provera (Pharmacia Upjohn, Kalamazoo, MI) subcutaneously to synchronize menstrual cycle and increase mouse susceptibility to *C. muridarum* infection.

### Monitoring vaginal *C. muridarum* burdens

To monitor vaginal live organism shedding, vaginal swabs were taken on different days after the intravaginal infection. As indicated in different data sets, slightly different schedules for collecting vaginal swabs were followed in different experiments. However, the overall schedule was once every week until two consecutive negative detection results were obtained from the same mouse. Most mice cleared infection by day 28 regardless of the immunization, which was why the chlamydial burdens from the 1^st^ month were used for comparison between groups. For determining the chlamydial burden in swabs, each swab was soaked in 0.5ml of SPG and vortexed with glass beads and the chlamydial organisms released into the supernatants were titrated on HeLa cell monolayers in duplicates as described previously (71). Briefly, the serially diluted swab samples were inoculated onto HeLa cell monolayers grown on coverslips in 24-well plates. After incubation for 24 hours in the presence of 2μg/ml cycloheximide, the cultures were processed for immunofluorescence assay (see below) and the inclusions were counted under a fluorescence microscope. Five random fields were counted per coverslip. For coverslips with less than one IFU per field, the entire coverslips were counted. Samples showing obvious cytotoxicity of HeLa cells were excluded. The number of IFUs per swab was calculated based on the number of IFUs per field, number of fields per coverslip, dilution factors and inoculation and total sample volumes. An average was taken from the serially diluted and duplicate samples for any given swab. The IFUs/swab were converted into log_10_ for calculating both chlamydial burden group means and standard deviation at each time point and area-under-curve (AUC) from each mouse and group comparison (see statistics below).

### Evaluating mouse genital tract pathology hydrosalpinx

Mice were sacrificed 50 to 60 days after infection for evaluating urogenital tract tissue pathology. Before removing the genital tract tissues from mice, an *in situ* gross examination was performed for evidence of hydrosalpinx formation and any other gross abnormalities. The severity of oviduct hydrosalpinx was semi-quantitatively scored as the following (71): oviduct with no hydrosalpinx was scored with 0; hydrosalpinx can only be seen after amplification under a stereoscope was scored with 1; Hydrosalpinx clearly visible with naked eyes but with a size smaller than that of the ovary on the same side was scored with 2; Hydrosalpinx with a size similar to that of ovary was scored with 3 while larger than the ovary scored with 4. The scores from both oviducts from the same mouse were added as the hydrosalpinx score of the mouse. The genital tissue gross pathology was documented with a digital camera.

### Immunofluorescence assay

HeLa cells grown on glass coverslips in 24 well plates were pretreated with DMEM containing 30mg/ml of DEAE-Dextran (Sigma, St Luis, MO) for 10 min. After the DEAE-Dextran solution was removed, the serially diluted swab samples were added to the monolayers and allowed to attach to the cell monolayers for 2 hours at 37°C. The infected cells were cultured in DMEM with 10% FCS and 2μg/ml of cycloheximide (Sigma) for 24h. The infected cultures were processed by fixing with 2% paraformaldehyde dissolved in PBS for 30 min at room temperature, followed by permeabilization with 2% saponin (Sigma) for 1h. After washing and blocking, the cell samples were labeled with Hoechst (blue, Sigma) for visualizing DNA and a rabbit anti-chlamydial organisms (unpublished data) plus a goat anti-rabbit IgG conjugated with Cy2 (green; Jackson ImmunoResearch Laboratories, Inc., West Grove, PA) for visualizing chlamydial inclusions. The immuno-labeled cell samples were quantitated as described above and used for image acquisition with an Olympus AX-70 fluorescence microscope equipped with multiple filter sets (Olympus, Melville, NY) as described previously (66, 72).

### Statistical analyses

For IFU data, the time course IFUs from each mouse were calculated into the area-under-curve (AUC) by adding up all measured IFUs and then dividing by the number of time points measured within the 1^st^ 4 weeks after the challenge infection. As a result, each mouse was rendered with one AUC value for group comparisons. For hydrosalpinx scores (HS) from each mouse, the scores from both oviducts were added up as the score for the mouse. When ANOVA Test (http://www.physics.csbsju.edu/stats/anova.html) was performed to analyze the IFUs (AUCs) and HS among multiple groups from a given immunization regimen, significant differences were found for both IFUs and HS. Then, Wilcoxon-signed rank test (2-tailed) was used to compare each antigen group with the corresponding negative and/or positive immunization control groups (http://faculty.vassar.edu/lowry/wilcoxon.html). The Fisher’s Exact test (http://www.danielsoper.com/statcalc/calc29.aspx) was used to compare incidence of positivity for infection or hydrosalpinx. Sine the analyses were carried out after all 17 independent experiments were completed, we were able to pool the controls from different experiments under the same regimen together for comparing with different antigen groups. Similarly, data for the same antigen repeated in different experiments under the same immunization regimen was also pooled for statistical analysis.

## Results

### 1. Intramuscular immunization with adjuvanted *C. muridarum* induced significant protection against subsequent chlamydial infection in the female mouse genital tract

Since intramuscular injection is a well-accepted route for vaccinating humans, per our industrial partner’s recommendation, we first evaluated whether the whole *C. muridarum* (CM) organisms without (live CM) or with UV-light inactivation (UV-CM) can induce protective immunity against subsequent challenge infection with *C. muridarum* in the genital tract. Because our goal is to establish a mouse model for evaluating protection efficacy of chlamydial antigens identified in humans (48), which requires adjuvants to enhance immunity, we used CpG and incomplete Freund’s adjuvant (IFA) to emulsify live CM or UV-CM. The adjuvanted CM preps were intramuscularly injected into mice three times and one month after the last immunization, mice were intravaginally challenged with live CM for evaluating the protection efficacy. As shown in Fig.1, all 30 mice that received intramuscular injection with adjuvanted live CM developed significantly reduced and shortened shedding courses of live chlamydial organisms from the genital tract (thus, designated as positive immunization controls) while 60 mice that received PBS with or without adjuvants or the adjuvanted control antigen glutathione-s-transferase (GST) displayed significantly high levels of live chlamydial organism shedding (collectively designated as mock or negative immunization controls). Interestingly, 15 mice that received adjuvanted UV-CM still developed moderate levels of genital shedding, which were significantly lower than those of the negative control groups but still higher than those of the positive immunization controls. These results were summarized from 8 independent experiments, which have demonstrated that the intramuscular immunization with CpG and IFA as adjuvants can be used to evaluate the protection efficacy of chlamydial antigens. It is not clear why the adjuvanted live CM induced significantly stronger protection against chlamydial infection in the genital tract than the adjuvanted UV-CM since the emulsification process is expected to inactivate all CM and muscle cells are considered non-permissible to chlamydial infection.

**Fig. 1.**
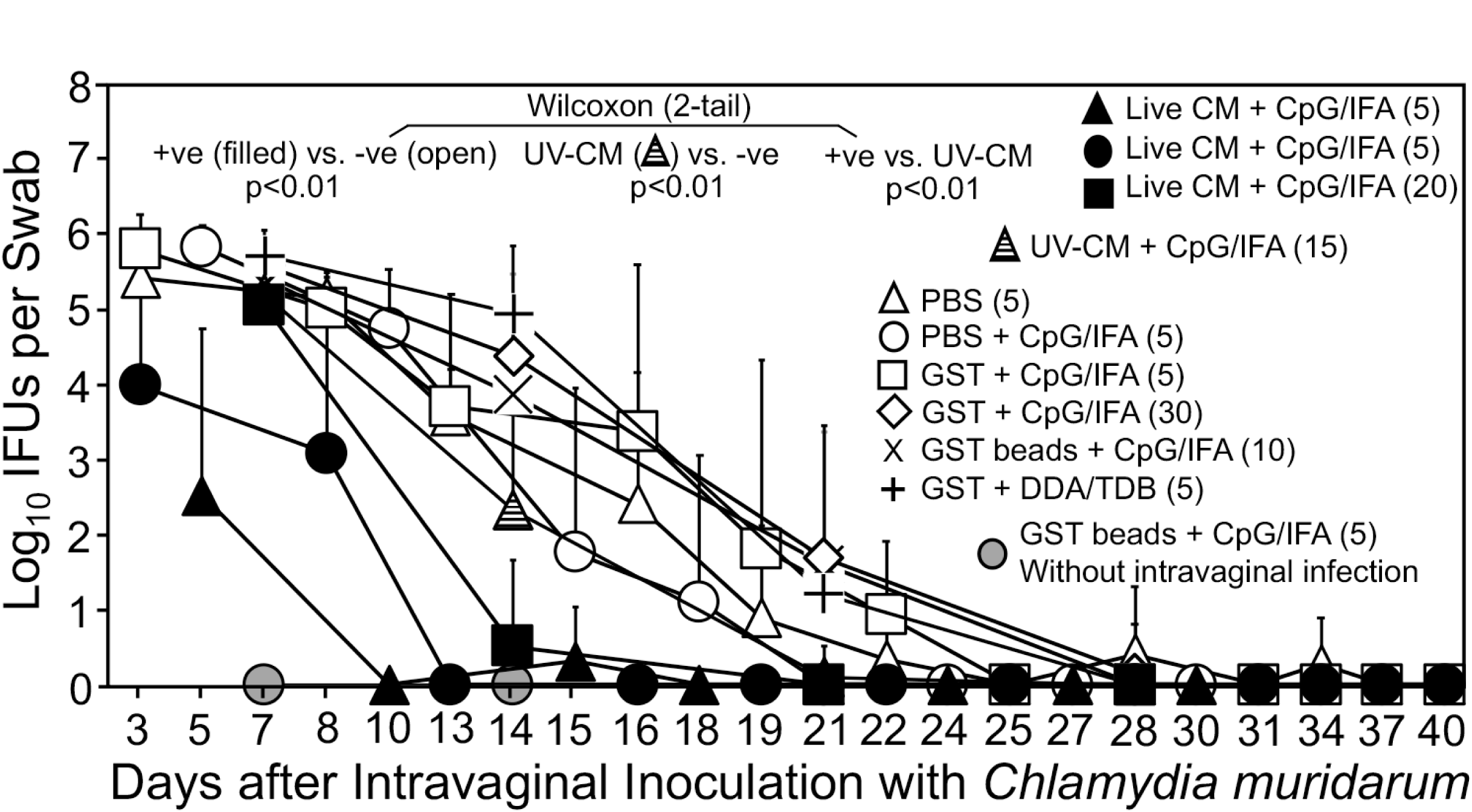
Chlamydia muridarum shedding courses from the genital tracts of mice intramuscularly immunized with adjuvanted *C. muridarum* or adjuvant alone. Groups of female mice were intramuscularly immunized three times on days 0, 20 & 30 respectively with UV light-inactivated *C. muridarum* (UV-CM, hatched triangle, n=15) or live CM (filled or solid symbols) adjuvanted with CpG and incomplete Freund’s adjuvant (IFA) or PBS buffer or GST (glutathione-s-transferase) with or without CpG/IFA (open symbols) or DDA/TDB adjuvants (+). For some groups, GST-bound to agarose beads were used as immunogens (X & grey circle). One month after the last immunization, all mice (except for the no infection control group, grey circle, n=5) were challenged intravaginally with 2×10^4^ IFUs (inclusion forming units) of *C. muridarum* organisms and vaginal swabs were taken once or twice weekly as indicated along the x-axis for monitoring genital chlamydial burdens. The vaginal swab collection from different groups started on different days, including days 3 (filled circle, n=5; open triangle, n=5; open square, n=5), 5 (filled triangle, n=5; open circle, n=5) or 7 (grey circle, n=5; filled square, n=20; open diamond, n=30; cross, n=10; plus, n=5) after intravaginal infection. The vaginal swabs were titrated for chlamydial burdens in terms of IFUs which were converted into Log10 for calculating group mean and standard deviation at a given time point as displayed along the y-axis. The IFU data were from 8 independent experiments. The live CM-immunized mice were regarded as positive immunization groups (+ve, filled symbols, n=30) while mice immunized with buffer or GST with or without adjuvants were considered negative immunization control groups (open symbols and cross/plus symbols, n=60). The area-under-curve (AUC) was compared between the +ve and -ve groups using Wilcoxon, p<0.01. The UV-EB group was different from both the +ve and -ve controls (p<0.01).

### 2. The intramuscular immunization regimen reveals more chlamydial antigens that induce detrimental responses and fewer antigens that induce protection

Using the established intramuscular immunization and adjuvant regimen, we evaluated 106 chlamydial antigens for protecting against the subsequent chlamydial infection in the female genital tract (Fig. 2). The native MOMP was purified from *C. muridarum* (73) while the remaining 105 antigens were purified from a bacterial expression system (48). When the vaginal chlamydial burdens of the antigen-immunized groups were compared with those of the negative immunization controls, two antigens were found to induce significant protection against genital chlamydial infection (thus designated as infection-reducing antigens) while surprisingly 11 were found to significantly exacerbate genital chlamydial infection (infection-enhancing antigens). The remaining 93 antigens did not significantly alter the live chlamydial shedding courses. The two infection-reducing antigens are native MOMP and TC0420. Native MOMP has been shown to induce significant protection (73-75) while TC0420 is a hypothetical protein that contains T cell epitopes (76) and has been shown to induce protective immunity against chlamydial infection (77). Thus, our screening system has validated previous observations. However, the same model system has also identified 11 infection-enhancing antigens. The finding that the intramuscular immunization regimen revealed more chlamydial antigens for enhancing/promoting than preventing chlamydial infection in the female mouse genital tract has not been documented previously.

**Fig. 2.**
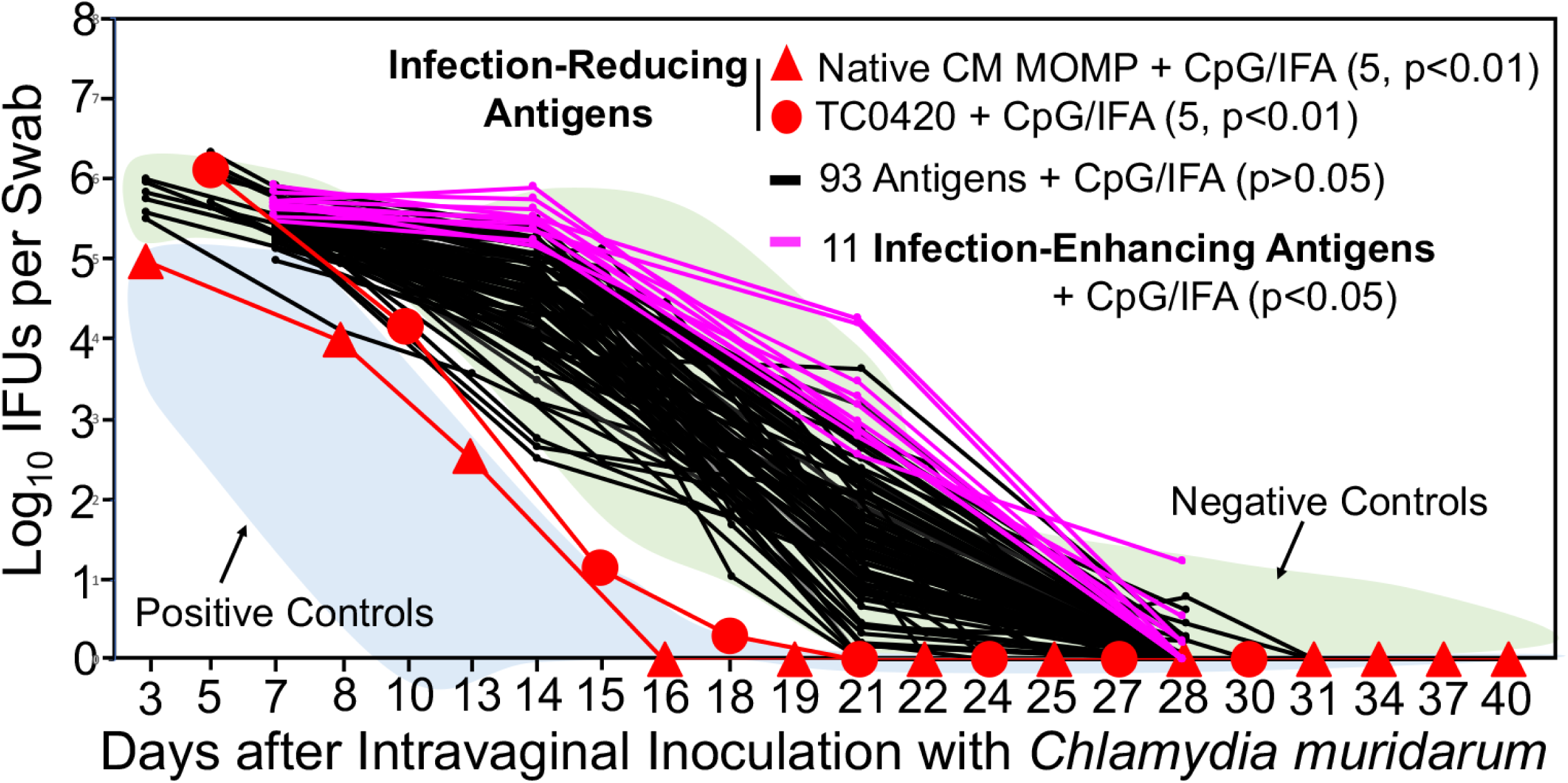
*Chlamydia muridarum* shedding courses from the genital tracts of mice intramuscularly immunized with different chlamydial antigens. Groups of female mice (n=4 to 10) were intramuscularly immunized with ∼30μg of each of 106 chlamydial antigens adjuvanted with CpG+IFA or DDA/TDB for 3 times for evaluating the intramuscular immunization-induced protection against genital tract infection as described in Fig.1 legend. The areas under curves of positive (+ve, light blue) and negative (-ve, light green) control groups from Fig.1 were shaded as references (indicated by arrows). A 2-tailed Wilcoxon analysis revealed that 2 antigens (native MOMP, filled red square; TC0420, filled red circle) significantly reduced genital chlamydial burdens (designated as infection-reducing antigens, red) while 11 antigens significantly enhanced chlamydial burdens (designated as infection-enhancing antigens, pink) when compared with the negative controls (-ve). Immunization with any of the remaining 93 antigens failed to significantly alter the chlamydial shedding courses (black curves). The IFU data were from the same 8 independent experiments as described in Fig.1 legend. To maintain clarity, the standard deviations were omitted from the plot.

When the upper genital tract pathology was compared between the negative immunization control groups and the chlamydial antigen-immunized groups (Fig. 3), no antigen groups, including the two infection-reducing antigen groups, significantly reduced hydrosalpinx scores. Surprisingly, 23 antigens significantly increased hydrosalpinx scores (designated as pathology-enhancing/exacerbating antigens; also see supplementary Fig.1S). Further, only one of the 23 pathology-exacerbating antigens significantly enhanced genital infection and the remaining 22 antigens did not, suggesting that the 22 antigens might have induced host responses for selectively promoting chlamydial pathogenicity without enhancing chlamydial infection. The identification of large number of pathology-enhancing antigens raised a concern on the safety of the intramuscular immunization regimen. As positive immunization controls, 30 mice that were intramuscularly immunized with live CM significantly prevented genital chlamydial infection (marked as +ve in Fig. 1). However, these same mice failed to consistently prevent pathology.

**Fig. 3.**
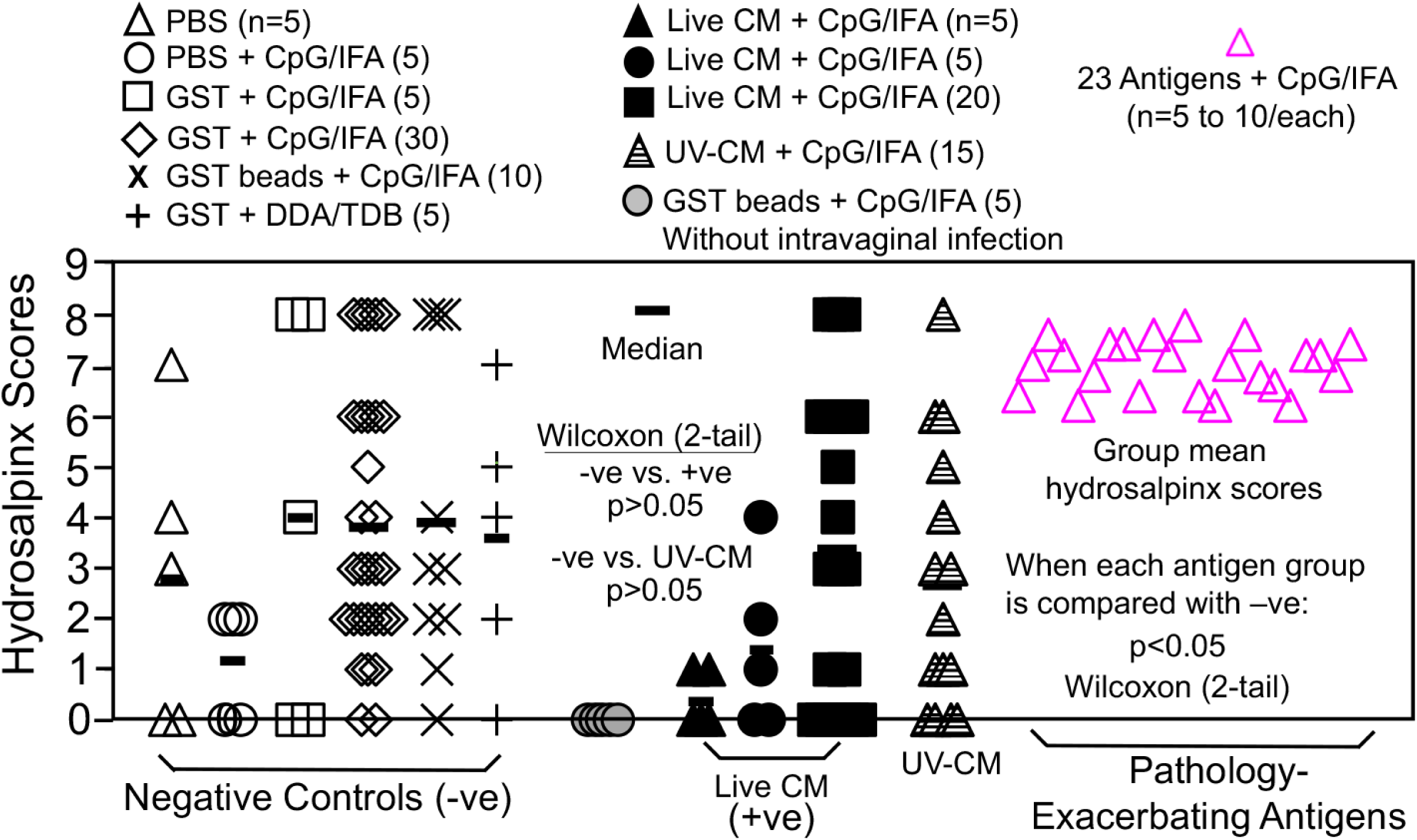
Hydrosalpinx scores of mice intramuscularly immunized with adjuvanted *C. muridarum* organisms or antigens or buffer/control antigen/adjuvant alone. Groups of female mice (n=4 to 10) were intramuscularly immunized and intravaginally challenged as described in Fig 1. legend. On day 56 after intravaginal challenge, all mice were sacrificed for semi-quantitatively scoring upper genital tract pathology hydrosalpinx as described in the materials and methods section and the hydrosalpinx scores were displayed along the Y-axis. A 2-tailed Wilcoxon analysis revealed no statistically significant difference in hydrosalpinx scores between the negative immunization controls (n=60, -ve) and positive controls (n=30, +ve) or UV-CM (n=15). Mice intramuscularly immunized with adjuvanted GST beads but without intravaginal challenge infection (grey circle) did not develop any significant pathology. When the hydrosalpinx scores from groups of mice (n=4 to 10) immunized with each of the 106 chlamydial antigens as described in Fig.2 legend were compared with that of the negative controls (n=60), 23 antigens were found to significantly increase hydrosalpinx (each pink open triangle represents the mean score from each of the 23 groups; for scores from individual mice in these 23 groups, please see supplementary Fig.1S). There was no significant difference in hydrosalpinx score between any of the remaining 83 antigen groups and the negative immunization control mice (data not shown). The hydrosalpinx score data were from the same 8 independent experiments as described in Figs.1 & 2 legends.

### 3. Intranasal mucosal immunization with live chlamydial organisms consistently prevents both chlamydial infection and pathogenicity in the female genital tract

To identify a more sensitive/appropriate immunization regimen for evaluating the protection efficacy of chlamydial vaccine antigen candidates, we next tested the intranasal mucosal immunization regimen. As shown in Fig. 4, all 26 mice each intranasally inoculated with a single dose of 50 or 500 IFUs of live CM developed immunity that significantly reduced genital chlamydial burden when compared with that of the 42 mice that were intranasally inoculated with PBS with or without GST and/or CpG adjuvant. Thus, the live CM-immunized mice were designated as positive immunization controls or +ve while mice receiving buffer or GST with or without CpG as negative immunization controls or -ve. It is worth noting that the 25 mice that were intranasally inoculated with CpG-adjuvanted UV-CM also significantly reduced vaginal chlamydial burden but their protection efficacy was significantly less potent than that induced by live CM. When the pathology was compared (Fig. 6), all live CM-immunized mice significantly prevented pathology while UV-CM-immunized mice failed to do so. Thus, intranasal inoculation with live CM but not UV-CM protected against both genital tract infection and pathology.

**Fig. 4.**
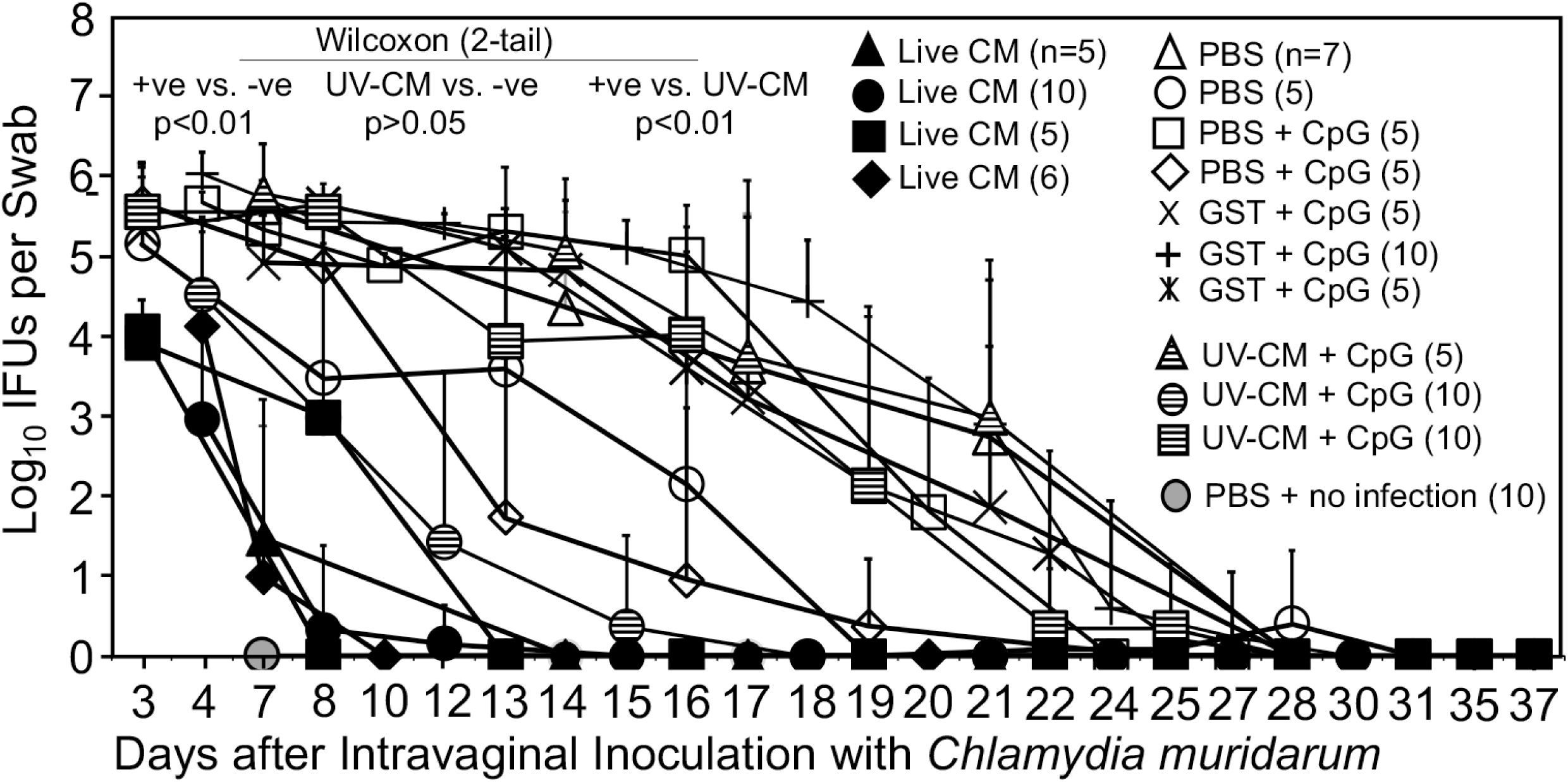
Chlamydia muridarum shedding courses from the genital tracts of mice intranasally immunized with *C. muridarum* or buffer/control antigen/adjuvant alone. Groups of mice were intranasally immunized once with live *C. muridarum* in PBS buffer (live CM, filled symbols) or immunized three times on days 0, 20 & 30 respectively with UV light-inactivated *C. muridarum* adjuvanted with CpG (UV-CM + CpG, hatched symbols) or PBS buffer or GST (glutathione-s-transferase) with or without CpG (open symbols). One month after the last immunization (2 months after live CM), all mice (except for the no infection control group, grey circle, n=5) were challenged intravaginally with 2×10^4^ IFUs of *C. muridarum* and vaginal swabs were taken once or twice weekly as indicated along the x-axis for measuring the number of live chlamydial organisms. The vaginal swab collection from different groups started on different days, including days 3 (filled triangle & square, n=5/each; open circle, n=5; hatched square, n=10), 5 (filled circle, n=10; filled diamond, n=6; open square, n=5; plus, n=10; hatched circle, n=10) or 7 (grey circle, n=10; cross, n=5; hatched triangle, n=5) after intravaginal infection. The vaginal swabs were titrated for chlamydial burdens in terms of IFUs which were converted into Log10 for calculating group mean and standard deviation at a given time point as displayed along the y-axis. The IFU data were from 5 independent experiments. The live CM-immunized mice were used as positive immunization controls (+ve, filled symbols, n=26) while mice immunized with buffer or GST with or without adjuvants as negative immunization controls (-ve, open symbols and cross/plus symbols, n=42). The area-under-curve (AUC) was used to compare between the +ve and -ve groups using Wilcoxon (p<0.01). The UV-CM groups (n=25) were significantly different from the +ve (p<0.01) but not the -ve (p>0.05) controls, indicating that intranasal inoculation with live CM but not killed CM significantly prevented genital chlamydial infection.

### 4. The intranasal immunization regimen fails to identify chlamydial antigens that can protect against both genital chlamydial infection and pathogenicity

We next used the intranasal immunization regimen for evaluating the protection efficacy of 29 chlamydial antigens (including antigens that were repeated multiple times such as various versions of Pgp3 and CPAF). Only two antigens were found to significantly reduce vaginal chlamydial burden (Fig. 5) and they were TC0511 and pCM03 or Pgp3. TC0511 is the recombination protein RecR with 203 amino acids (Genbank#AAF39353). Although TC0511 is a highly conserved chlamydial protein with significant homology with its bacterial homologues, its homologue (CT240) in *C. trachomatis* does not have any significant homology with any human proteins based on blastp search. Thus, CT240 could be considered a human vaccine candidate if it were also able to prevent pathology. The 2^nd^ infection-reducing antigen is the most immunodominant *C. muridarum* plasmid-encoded glycoprotein Pgp3 (78). A total of 35 mice in 5 different groups with 5 to 10 mice per group were intranasally immunized with Pgp3 in 4 independent experiments (Table 1S). It is worth emphasizing that intranasal immunization with Pgp3 was found to significantly reduce genital chlamydial burden only in one of the 4 experiments. The remaining Pgp3-immunized 30 mice failed to significantly reduce the burden. When all Pgp3-immunized 35 mice were compared with the 42 negative control mice, no significant difference in genital chlamydial burden was found. These results have demonstrated that pCM03 may not be considered a protective antigen. Moreover, when the pathology was compared (Fig. 6), only one group of 10 mice immunized with Pgp3 significantly reduced the hydrosalpinx score but without impacting the genital chlamydial burden. Paradoxically, another group of 10 mice intranasally immunized with Pgp3 were found to significantly increase the hydrosalpinx score but without enhancing genital chlamydial infection. When the hydrosalpinx scores of all Pgp3-immunized mice were compared with those of the negative control mice, no more significant difference was found. Thus, Pgp3 (without modification) may not be a reliable vaccine antigen under the intranasal immunization regimen. Further, Fig. 6 also revealed two additional pathology-exacerbating antigens: TC0043 and TC0911. TC0043 is the chlamydial type III secretion translocase SctQ with 373 amino acids (Genbank#AAF73521) while TC0911 is a conserved hypothetical protein with 831 amino acids (Genbank#AAF39704; ref: (60)). Intranasal inoculation with TC0043 or TC0911 significantly exacerbated oviduct hydrosalpinx but without significantly altering the genital infection, suggesting that these two antigens-induced immune responses might directly promote pathogenicity in the upper genital tract. It will be interesting to further characterize the pathology-exacerbating responses. Overall, the intranasal immunization regimen failed to identify any chlamydial antigens that can prevent both genital chlamydial infection and pathology, suggesting that more effective immunization regimen is required for identifying chlamydial vaccine antigens.

**Fig. 5.**
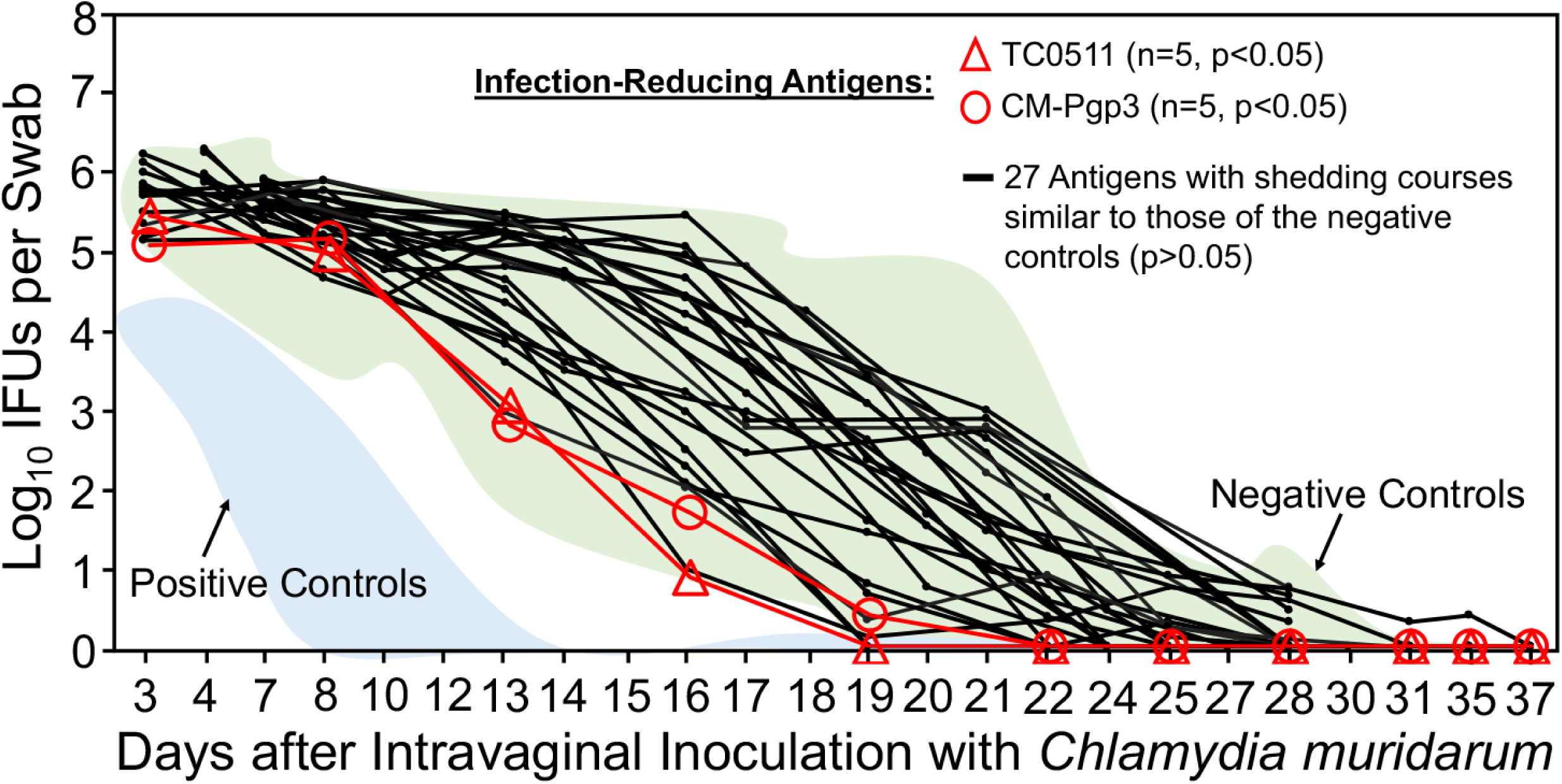
*Chlamydia muridarum* shedding courses from the genital tracts of mice intranasally immunized with different chlamydial antigens. Groups of female mice (n=5 to 10) were intranasally immunized with ∼30μg of each of 29 chlamydial antigens adjuvanted with CpG for 3 times for evaluating the intranasal immunization-induced protection against genital tract infection as described in Fig.4 legend. The areas under curves of positive (+ve, light blue) and negative (-ve, light green) control groups from Fig.4 were shaded as references (indicated by arrows). A 2-tailed Wilcoxon revealed that mice immunized with TC0511 (red open triangle) or CM-Pgp3 (red open circle) significantly reduced genital chlamydial burdens (designated as infection-reducing antigens) when compared with the negative controls (-ve). Immunization with any of the remaining 27 antigens failed to significantly alter the chlamydial shedding courses (black). The IFU data were from the same 5 independent experiments as described in Fig.4 legend. To maintain the presentation clarity, the error bars were not omitted.

**Fig. 6.**
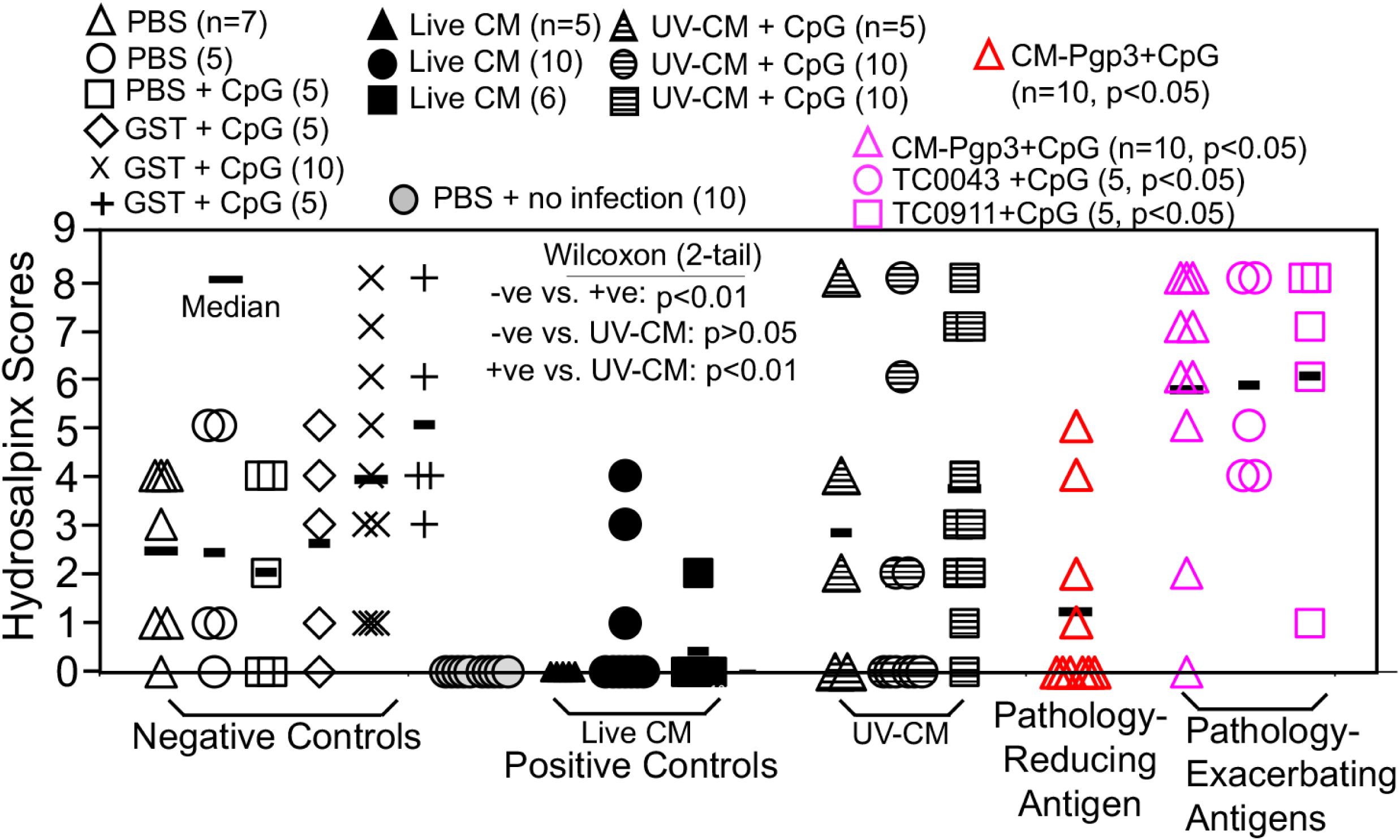
Hydrosalpinx scores of mice intranasally immunized with *C. muridarum* organisms or antigens or buffer/control antigen/adjuvant alone. Groups of mice were intranasally immunized and intravaginally challenged as described in Figs. 4 & 5 legends. On day 56 after intravaginal challenge, all mice were sacrificed for semi-quantitatively scoring the upper genital tract pathology hydrosalpinx as described in the materials and methods section and the hydrosalpinx scores were displayed along the Y-axis. A 2-tailed Wilcoxon analysis revealed statistically significant difference in hydrosalpinx scores between the negative immunization controls (-ve, open, n=37) and positive controls (+ve, filled, n=21, p<0.01) but not the UV-CM (hatched symbols, n=25, p>0.05). Mice intranasally immunized with PBS but without intravaginal challenge infection (grey circle) did not develop any significant pathology. The hydrosalpinx score data were from the same 5 independent experiments as described in Fig.4 & 5 legends except for groups from VacExp B, the pathology data were missing (see supplementary Table 1S). When each of the 29 chlamydial antigen groups (n=4 to 10) was compared with the negative controls (n=37), the plasmid-encoded Pgp3 significantly reduced the hydrosalpinx score (red open triangle) while the same Pgp3 in a different experiment (pink open triangle) and antigens TC0043 (pink open circle) & TC0911 (pink open square) significantly increased the hydrosalpinx scores. Note that intranasal immunization with live CM but not UV-CM significantly prevented genital pathology while CpG-adjuvanted chlamydial antigens either reduced or exacerbated pathology in different experiments. Particularly, Pgp3 was repeated 5 times with a total of 35 mice under the same intranasal immunization regimen (see supplementary Table 1S) and the hydrosalpinx scores of the 35 mice were not significantly different from those of the negative control mice.

### 5. A combinational immunization regimen identifies a chlamydial antigen that prevents both chlamydial infection and pathology in the female genital tract

Since neither the intramuscular nor the intranasal immunization regimen has identified chlamydial antigens that can prevent both chlamydial infection and pathology, we next tested a combinational immunization regimen consisting of intranasal priming, intramuscular boosting with or without a 3rd boost intraperitoneally or subcutaneously. As shown in Fig. 7, all 30 mice receiving intranasal priming with live CM and intramuscular boosting with adjuvanted UV-CM with or without the 3^rd^ booster significantly reduced the shedding of live chlamydial organisms comparing to the negative control mice. When groups of mice immunized with individual chlamydial antigens were compared with the negative control mice, 4 antigens were found to significantly reduce genital chlamydial infection: TC0151, TC0453, TC0052c-Pgp3c fusion protein, and TC0727c. TC0151 is beta-ketoacyl-[acyl-carrier-protein] synthase II (fabF, WP_010229532.1) while TC0453 is a hypothetical protein (WP_010230489.1). TC0052 is the major outer membrane protein (MOMP) while TC0727 is the outer membrane complex protein B (OmcB). The rationale of testing the TC0052c-Pgp3c fusion protein was to take advantage of the Pgp3 C-terminal trimerization domain to trimerize the C-terminal half of MOMP that contains variable domains (VD) III and IV (79). The rationale of testing the C-terminus of OmcB was that the immunogenicity of OmcB in humans and chlamydia-infected animals was largely mapped to its C-terminus (80, 81). We also found that TC0171 (a DUF4339 domain-containing protein, WP_010229701.1) and Pgp3 significantly enhanced chlamydial infection under the same combinational immunization regimen, which once again suggested that Pgp3 is not a safe vaccine antigen. When the pathology was compared (Fig. 8), all live CM-primed mice were prevented from any significant pathology while significant pathology developed in the negative control groups (p<0.01). Mice immunized with chlamydial antigens TC0043 (SctQ, WP_010229202.1), TC0079 (ATP-dependent Clp protease proteolytic subunit, WP_010229313.1), TC0151 (beta-ketoacyl-ACP synthase II, fabF, WP_010229532.1), TC0375 (thioredoxin-disulfide reductase, trxB, WP_010230279.1) or TC0892 (peroxiredoxin, WP_010231862.1) all significantly reduced pathology. Among the 5 pathology-reducing antigens, only TC0151 significantly prevented chlamydial infection in the genital tract as shown in Fig. 7. Thus, we have designated TC0151 as a protective chlamydial antigen. Finally, Fig. 8 also showed that mice immunized with TC0066 or TC0376 significantly enhanced pathology although these two antigens failed to significantly promote chlamydial infection. It is worth noting that although TC0043 reduced pathology under the combinational immunization regimen, the same antigen exacerbated pathology under the intranasal immunization regimen although TC0043 did not affect chlamydial challenge infection courses in the genital tract under either regimen. These observations suggest that the immunization routes or adjuvants could affect the immune phenotypes that can either reduce or exacerbate pathology. Despite the variable results from the individual antigen-immunized mice, all live CM-primed mice (regardless of the subsequent boosts) consistently prevented both genital chlamydial infection and pathology, suggesting that the variations among chlamydial antigen-immunized mice were not caused by the mouse model system. On the contrary, the findings from the mouse model studies have consistently demonstrated the importance of priming mice with live chlamydial organisms in inducing adequate and appropriate immune responses for preventing both genital infection and pathology.

**Fig. 7.**
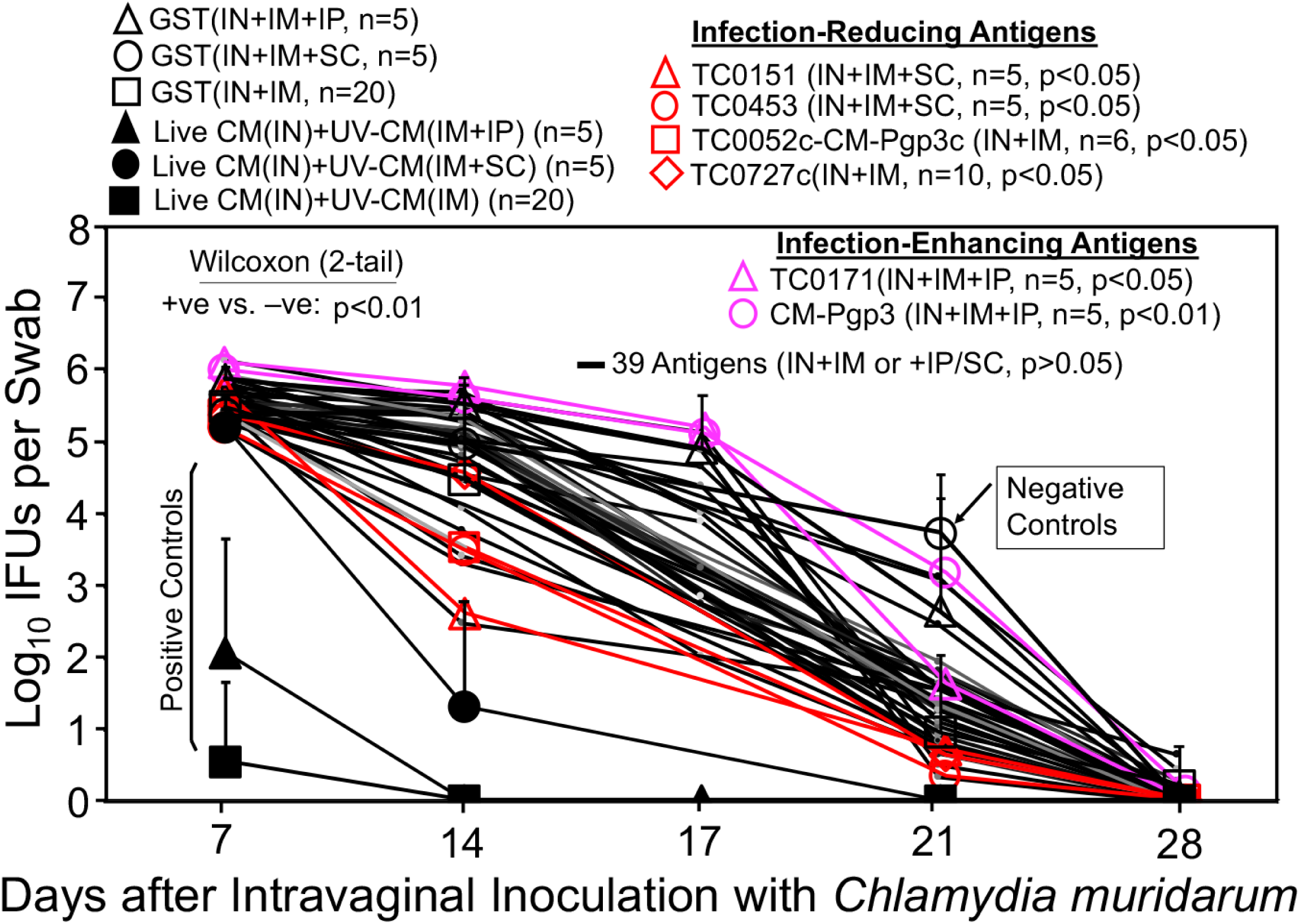
Chlamydia muridarum shedding courses from the genital tracts of mice immunized via a combinational regimen. Groups of mice were immunized three times with intranasal (IN) priming on day 0, intramuscular (IM) boosting on day 20 and a final intraperitoneal (IP) or subcutaneous (SC) booster on day 30 respectively with *C. muridarum (*filled symbols as positive immunization controls, +ve), GST (open symbols as negative immunization controls, -ve) or different chlamydial antigens (red & pink symbols and black lines). One month after the last immunization, all mice were challenged intravaginally with 2×10^4^ IFUs of *C. muridarum* and vaginal swabs were taken once weekly as indicated along the x-axis for monitoring chlamydial burdens in terms of Log10 IFUs as displayed along the y-axis. The area-under-curves (AUCs) were compared between the +ve and -ve groups using a 2-tailed Wilcoxon (p<0.01). Comparing to the -ve controls, chlamydial antigens TC0151, TC0453, TC0052c-Pgp3c, and TC0727c were found to significantly reduce genital chlamydial burdens (designated as infection-reducing antigens, red) while antigens TC0171 and Pgp3 significantly enhanced chlamydial burdens (designated as infection-enhancing antigens, pink). The IFU data were from the 4 independent experiments.

**Fig. 8.**
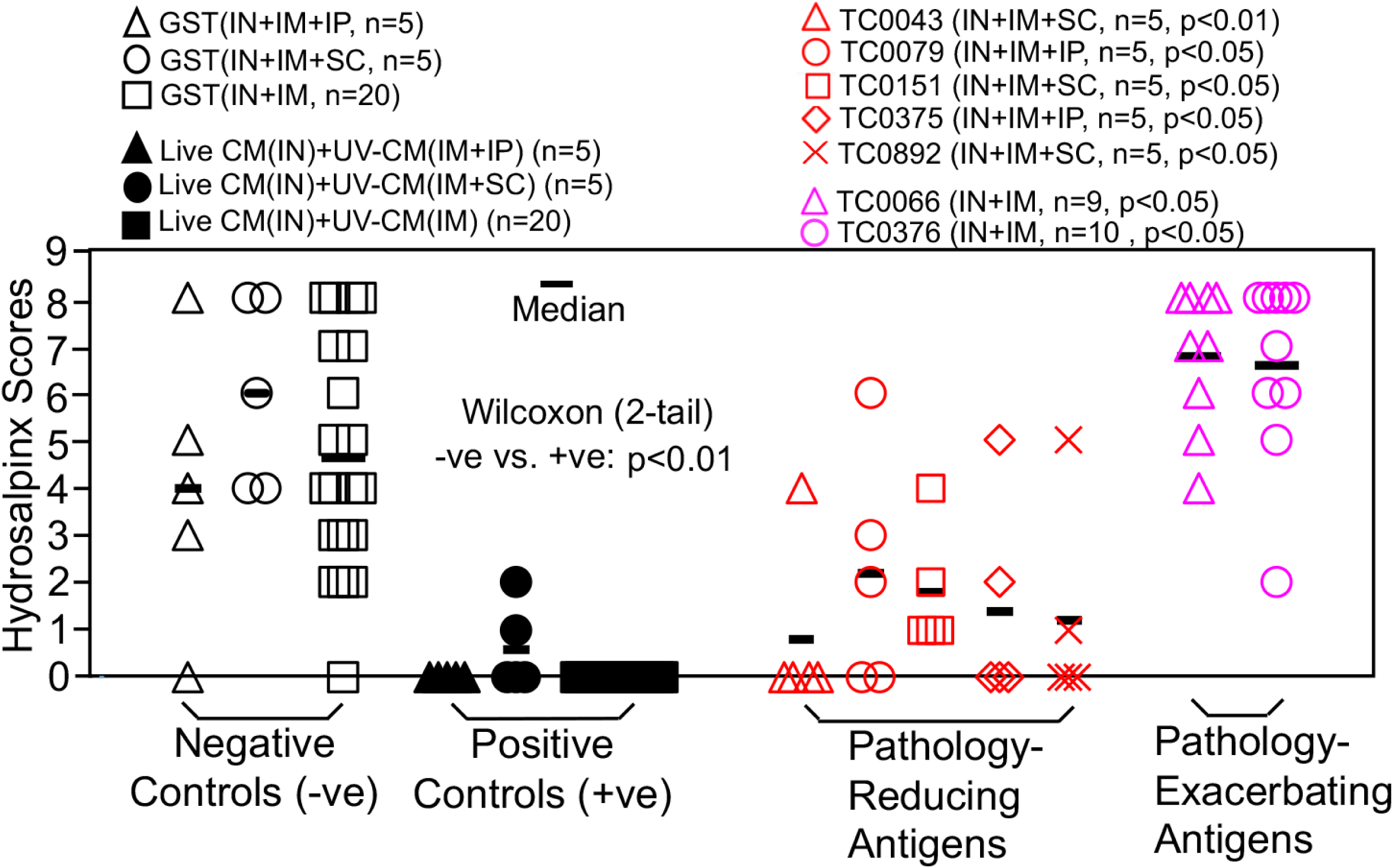
Hydrosalpinx scores of mice immunized via a combinational regimen. Groups of female mice (n=4 to 10) were immunized via a combinational regimen and intravaginally challenged as described in Fig. 7 legend. On day 56 after intravaginal challenge, all mice were sacrificed for semi-quantitatively scoring the upper genital tract pathology hydrosalpinx as described in the materials and methods section and the hydrosalpinx scores were displayed along the Y-axis. A 2-tailed Wilcoxon analysis revealed statistically significant difference in hydrosalpinx scores between the negative immunization controls (-ve, open, n=30) and positive controls (+ve, filled, n=30, p<0.01). Comparing to the -ve controls, chlamydial antigens TC0043, TC0079, TC0151, TC0375 and TC0892 were found to significantly reduce hydrosalpinx (designated as pathology-reducing antigens, red) while antigens TC0066 and TC0376 significantly enhanced hydrosalpinx (designated as pathology-enhancing antigens, pink). The hydrosalpinx score data were from the same 4 independent experiments as shown in Fig. 7. Note that only TC0151 prevented both genital chlamydial infection and pathology.

## Discussion

The successful isolation of *C. trachomatis* from human ocular tissues by Tang et al (82, 83) more than sixty years ago had laid the foundation for developing a Chlamydia vaccine. Since then, extensive efforts have been made for achieving the goal (21-38). However, no licensed *C. trachomatis* vaccine is available today. The current study was designed to search for chlamydial subunit vaccine antigens by screening 140 *C. trachomatis* antigens identified to be immunogenic in women infected with *C. trachomatis* (48) using a *C. muridarum* infection of female mouse genital tract model (41). The *C. muridarum* homologues (60) were used as immunogens in most cases to match the genital challenging infection organisms. The advantage of this mouse model is that it can evaluate vaccine efficacy by simultaneously monitoring both genital infection and pathology (42, 47, 84).

We started with the intramuscular priming plus two boosts regimen with CpG and IFA as adjuvants since intramuscular injection is a well-accepted route for vaccinating humans and CpG and IFA can promote both antibody and Th1 responses. After screening 106 chlamydial antigens, we did not find any chlamydial antigen that can prevent both chlamydial infection and pathology in the female mouse genital tract. Instead, we found that only two antigens reduced but 11 promoted chlamydial infection and no antigen prevented genital pathology but 23 exacerbated pathology. Although the failed human trachoma vaccine trials based on intramuscular immunization with formalin-fixed chlamydial organisms (21-24) had already suggested that chlamydial organisms might contain antigens that can induce detrimental immune responses for exacerbating chlamydial infection/pathogenicity, it was still surprising to find so many chlamydial antigens that enhanced chlamydial infection/pathogenicity in the mouse model. This phenomenon was rarely reported in previous mouse model-based chlamydial vaccine studies, which was probably because most studies were only focused on one or a few chlamydial antigens. More importantly, although the positive immunization control mice (mice immunized with adjuvanted live CM) reduced genital infection in multiple independent experiments, these mice were not always protected from pathology. None of the UV-CM-immunized mice was protected from pathology under the intramuscular immunization regimen. Thus, we tested the intranasal mucosal immunization regimen since intranasal inoculation with *C. muridarum* is known to prevent both genital tract infection and pathology (50). After evaluating 29 chlamydial antigens, we identified two infection-reducing antigens but no infection-enhancing antigens, which is an improvement over the intramuscular immunization regimen. However, neither of the infection-reducing antigens significantly prevented the genital tract pathology, suggesting that the prevention of pathology might require stronger immunity. Thus, we further tested a combinational immunization regimen with intranasal mucosal priming followed by intramuscular boosting with or without additional intraperitoneal or subcutaneous boosters. Among the 44 antigens evaluated, we found 4 infection-reducing antigens, two infection-enhancing antigens, 5 pathology-reducing antigens and two pathology-enhancing antigens. Clearly, the combinational regimen identified more protective antigens than detrimental antigens at both the anti-infection and anti-pathology levels. More importantly, TC0151 was found to prevent both genital tract infection and pathology. It is worth noting that all mice intranasally primed with live CM were protected from both genital chlamydial infection and pathogenicity. The subsequent boosting with intramuscular injection of adjuvanted UV-CM did not alter the protective phenotype of the immune responses, suggesting that mucosal priming with live CM is important for maintaining the protective phenotype of the induced immune responses.

Combinational immunization with TC0151 prevented both genital tract infection and pathology. TC0151 is a highly conserved beta-ketoacyl-[acyl-carrier-protein] synthase II (fabF, WP_010229532.1) with 418 amino acids that shares 98% amino acid identity with the *C. trachomatis* fabF (WP_009872915.1). FabF is involved in the condensation step of fatty acid biosynthesis in which the malonyl donor group is decarboxylated and the resulting carbanion used to attack and extend the acyl group attached to the acyl carrier protein. *C. trachomatis* fabF is immunogenic in *C. trachomatis*-infected women (48), suggesting that fabF-based vaccine-primed immune responses can be recalled during *C. trachomatis* infection in women. However, *C. trachomatis* fabF was found to have significant levels of amino acid identity/similarity with the various isoforms of human fatty acid synthase (for example, 24.5% identity against AAH63242.1) and human mitochondrial beta-ketoacyl ACP synthase (47.78% against XP_006713280.1). Careful comparison has revealed identical or similar amino acids (without gaps) in ∼10 different regions (each with 5 to 10 amino acids in length) between *C. trachomatis* fabF and its human homologues, raising a serious safety concern (on potentially inducing auto-reactive immune responses in humans when a *C. trachomatis* fabF-based vaccine is applied). Thus, although chlamydial fabF protected against both chlamydial infection and pathology in the female mouse genital tract, it was recommended not to move forward to the next step.

Some chlamydial antigens were found to significantly reduce genital tract infection but without reducing pathology while others significantly prevented pathology but without preventing genital infection. These antigens were not recommended to move forward for vaccine development. This is because an effective vaccine should prevent both infection and pathology. The discrepancy displayed by these antigens might be caused by either the inability of an antigen to induce adequate protective response or the inappropriate immunization regimens and adjuvants. The *C. muridarum* genital infection mouse model is considered sensitive and specific for measuring the protection efficacy against both genital infection and pathology. This conclusion is supported by the identification of chlamydial fabF as a protective antigen for preventing both genital chlamydial infection and pathology reported in the current study. The finding that mice intranasally primed with live CM were always protected against both genital chlamydial infection and pathology has further demonstrated the reproducibility of the mouse model. Thus, if a chlamydial antigen fails to protect against both genital infection and pathology in the mouse model, it is unlikely that this antigen will be successful in large animal models. Obviously, an effective subunit vaccine may combine multiple infection-reducing antigens with pathology-reducing antigens to form multi-valent vaccines. The antigen combination idea was tested in the current study by genetically fusing Pgp3 fragments with MOMPc or fragments of OmcB with fragments of CPAF in various versions. Only the fusion protein TC0052c-pCM03c was found to significantly reduce genital chlamydial infection but without preventing pathology. Although we still have plans to re-evaluate the protection efficacy of different antigen combinations, we are also aware of the safety concern that antigen combinations might also increase the risk of promoting infection and/or exacerbating pathology.

Many chlamydial antigens were found to induce detrimental responses for promoting genital infection and/or exacerbating pathology. The failed human trachoma vaccine trials had suggested that some chlamydial antigens might induce detrimental responses for exacerbating mucosal pathology upon chlamydial infection (21-23). Although the chlamydial antigens responsible for detrimental responses in humans and mice are not necessary the same, it will be interesting to investigate the mechanisms by which some of the antigens promote genital chlamydial infection and/or pathogenicity. It is worth pointing out that the chlamydial antigen-induced detrimental immune responses have not been widely reported using mouse models, but on the other hand, to the best of our knowledge no large-scale antigen screen has been performed for any chlamydia. Most detrimental chlamydial antigens were identified under the intramuscular immunization regimen, which is consistent with the observation made in the failed human trachoma vaccine trials since the trachoma vaccines were also based on intramuscular immunization with formalin-fixed whole chlamydial organisms. The intranasal immunization regimen enabled more chlamydial antigens to induce protective responses in the female mouse genital tract. Consistently, all mice intranasally immunized with live CM prevented both genital chlamydial infection and pathology. It will be interesting to investigate the mechanisms by which intranasal immunization with chlamydial antigens tend to induce protective responses.

A consistent finding in the current study is that immunization with live CM is always more effective than UV-CM in inducing protection against genital chlamydial infection and pathology. Even under the intramuscular immunization regimen, adjuvanted live CM induced significantly stronger protection against genital chlamydial infection than adjuvanted UV-CM did. Although emulsification with adjuvants is expected to inactivate CM and muscle cells are considered non-permissible to chlamydial infection, it is possible that some viable CM organisms may remain after emulsification. The remaining viable CM may still transiently infect mouse muscle cells and produce infection-dependent antigens for directly activating protective responses. Nevertheless, although intramuscular injection with adjuvanted live CM prevented genital tract chlamydial infection, it failed to consistently prevent pathology. Only after intranasal inoculation, was live CM able to fully prevent both genital chlamydial infection and pathology. This may be because the intranasally inoculated live CM can undergo productive infection in the airway, triggering strong protective immune responses. Consistently, live chlamydial infection-dependent protective immunity has been frequently observed (73, 85-88). The protective immunity induced by live CM was corelated with a Th1-dominant but Th17-low T cell response coupled with immune responses to chlamydial secretion proteins (50). Immune responses with similar phenotypes are thought to be important for clearing chlamydial infection (43). The challenge is how to induce the infection-dependent protective immune responses using non-infectious chlamydial antigen platforms. Some have tested surrogate live vectors such as adenovirus (89), poliovirus (90) or influenza viruses (91) while others have evaluated non-viable vectors such as Vibrio cholerae ghosts (VCG; ref: (26, 27)) and the immunogenic bacteriophage MS2 virus-like particle (VLP; ref: (92) for enhancing chlamydial antigen immunogenicity. These vector systems may function as both adjuvants and presentation platforms. For example, VCG was found to exert immunomodulatory effect on dendritic cells for enhanced antigen presentation and induction of protective immunity (93). A recent study has evaluated the protection efficacy of UV-inactivated EBs after modification with nanoparticles and pattern-recognition receptor ligands (34). However, none of these has been advanced to the stage of clinical evaluation. A most advanced chlamydia vaccine is CTH522 adjuvanted with CAS01 liposomes or aluminum hydroxide, for which the phase I trial was completed in 2019 (35).

Both the difficulty in developing subunit Chlamydia vaccines (29, 33) and the acquisition of genital tract pathogenicity-attenuated chlamydial mutants (54, 56, 58) have reignited the interest in developing whole chlamydial organism-based vaccines. In particular, the discovery that chlamydial colonization in the gastrointestinal tract is non-pathological and can induce transmucosal protection against subsequent chlamydial infection in the female genital tract has laid the foundation for developing an live-attenuated oral chlamydial vaccine (57). Consistently, the current study has demonstrated that mice with intranasal mucosal priming with live CM significantly prevented both genital chlamydial infection and pathogenicity but similar priming with UV-CM failed to do so. Oral immunization with various genital pathogenicity-attenuated CM mutants has been shown to induce protective immunity against both chlamydial infection and pathology in the female genital tract (54, 56, 58). Efforts are underway to move some of the attenuated mutants to clinical evaluation. The efforts are supported by the facts that live-attenuated *C. psittaci* (94) & *C. abortus* (95) vaccines are approved for protecting animals. The question is whether oral CM-induced immunity can cross-protect humans against *C. trachomatis* infection and pathology in the genital tract? First, there are successful examples using animal-adapted microbes as vaccines for protecting humans against human pathogens, including cowpox virus as a live vaccine to protect humans against smallpox (53), BCG of *M. bovis* against *M. tuberculosis* (51, 96) and meningococcal vaccine against gonorrhea (97). Second, prior exposure to CM has been reported to induce protective immunity against genital infection by multiple serovars of the human pathogen *C. trachomatis* (98). Thus, the next steps are to test whether oral delivery of an attenuated CM can protect against *C. trachomatis* in the genital tracts of large animal models or humans.

## Acknowledgements

This work was supported in part by grants (to G. Zhong) from Merck and NIAID.

## Figure legends and Table notes

**Supplementary Fig.1S: Hydrosalpinx scores from the negative immunization control mice and mice intramuscularly immunized with each of 23 chlamydial antigens.**

The hydrosalpinx scores from the 60 negative immunization control mice from Fig.3 were shown in black open triangles here and those from mice intramuscularly immunized with each of 23 chlamydial antigens listed along the X-axis were shown in pink open triangles. Note that hydrosalpinx scores from each of the 23 antigen groups were significantly higher than that of the negative immunization control mice (2-tailed Wilcoxon, p<0.05).

**Supplementary Table 1S: Vaginal chlamydial burdens and hydrosalpinx scores from antigen groups reported in the current manuscript.**

The data are organized into three different immunization regimens as listed in the 1^st^ column, including intramuscular (top of Table 1S), intranasal (middle) and combinational (bottom). The corresponding adjuvants used for each group are listed in the 2^nd^ column. CpG plus incomplete Freund’s adjuvant (IFA) were used for intramuscular, intraperitoneal, and subcutaneous injections while CpG alone for intranasal inoculation with the exception that no adjuvant was used for intranasal inoculation with live chlamydial organisms. The sample size for each group is listed in the 3^rd^ column. Mice with similar experimental conditions are combined into one group while those with different experimental conditions are kept in the original groups even though they were immunized with the same antigens. The group IDs and antigens used for immunizing the corresponding groups are listed in the 4^th^ column, which are from 17 different experiments as listed in the 7^th^ column or last column. All negative (green) and positive (blue) immunization controls from the same immunization regimen are clustered on top of each immunization regimen. The negative controls were used to compare with individual antigen groups under the same regimen to identify antigen groups with statistically different chlamydial burden (5^th^column) and hydrosalpinx score ( 6^th^ column). The chlamydial burden is listed as area-under-curves from each group. After ANNOVA analysis, a 2-tailed Wilcoxon Rank Sum test was used to do the comparison, which revealed both infection/pathology-reducing (red) and -enhancing/exacerbating (pink) antigens.

